# Coronavirus surveillance of wildlife in the Lao People’s Democratic Republic detects viral RNA in rodents

**DOI:** 10.1101/2020.04.22.056218

**Authors:** David J. McIver, Soubanh Silithammavong, Watthana Theppangna, Amethyst Gillis, Bounlom Douangngeun, Kongsy Khammavong, Sinpakone Singhalath, Veasna Duong, Philippe Buchy, Sarah H. Olson, Lucy Keatts, Amanda E. Fine, Zoe Greatorex, Martin Gilbert, Matthew LeBreton, Karen Saylors, Damien O. Joly, Edward M. Rubin, Christian E. Lange

## Abstract

Coronaviruses can become zoonotic as in the case of COVID-19, and hunting, sale, and consumption of wild animals in Southeast Asia facilitates an increased risk for such incidents. We sampled and tested rodents (851) and other mammals, and found *Betacoronavirus* RNA in 12 rodents. The sequences belong to two separate genetic clusters, and relate closely to known rodent coronaviruses detected in the region, and distantly to human coronaviruses OC43 and HKU1. Considering close human-wildlife contact with many species in and beyond the region, a better understanding of virus diversity is urgently needed for the mitigation of future risks.

## Brief Report

The latest coronavirus (CoV) outbreak in humans, caused by the SARS-CoV-2 virus [1], originated in Wuhan, Hubei Province, in the People’s Republic of China in late 2019. The suspected index case contracted the virus at a local seafood and wildlife market in the city, yet the exact species of animal that hosted the virus remains unknown. Phylogenetic analysis of the SARS-CoV-2 genome indicates a strong likelihood that the reservoir species is a bat, as in the case of the related *Betacoronaviruses* SARS-CoV-1 and MERS-CoV [2]. An involvement of another intermediate host between bats and humans in the transmission of SARS-CoV-2 virus remains unknown. Wildlife and bushmeat markets are common across Southeast Asia, and represent a significant risk for the transfer of zoonotic pathogens between wildlife and humans. Indeed, 75% of all emerging infectious diseases in the past decades have their origin in wildlife, including highly pathogenic influenza viruses (H5N1), Ebolaviruses, Henipaviruses, Hantaviruses among others [3].

The CoVs most closely related to SARS-CoV-2 were isolated from bats living in Yunnan province, in the south of China, not far from the 423km long border with the landlocked Lao People’s Democratic Republic (Laos) [4]. Both countries were involved in the United States Agency for International Development’s (USAID) Emerging Pandemic Threats PREDICT program, and surveillance of bats in wildlife markets in rural areas in Laos unveiled CoV RNA in 41 animals using family-level PCR assays [5]. In addition to bats, rodents are recognized as significant hosts of viral zoonoses, and represent an important potential host for zoonotic viral spill over in Laos, through their frequent incidental and intentional interaction with humans [6,7].

Multiple groups in Laos are at high risk of zoonotic viral spillover from wildlife, including from rodents, due to their occupation, economic or geographic circumstances. People contact rodents incidentally and intentionally in various ways. In traditional-style homes, especially in rural areas, rodents are often able to easily enter the houses in search of food and shelter. These circumstances promote incidental contact with rodent urine and feces during everyday life, when household members clean their houses. Rodents also commonly raid food storage areas, including rice storage huts near paddies. In terms of more direct and intentional contact, some species of rodents, including the Indian giant flying squirrel (*Petaurista philippensis*), Finlayson’s squirrel (*Callosciurus finlaysonii*), red-cheeked flying squirrel (*Hylopetes spadiceus*), and others, are hunted or trapped in rural forested areas using traps, guns, sticks, or other implements. Designated for food or for medicinal purposes, depending on the species, the rodents are consumed within the hunter’s village, or enter the value chain to reach markets. The value chain involves a series of intermediaries that transit animals from small villages to progressively larger populated areas. At the market, animals are sold to locals or to Lao people visiting from other areas of the country, and often to foreign visitors from neighbouring Thailand, China, and Vietnam [8,9]. Even though the sales of these animals are illegal, Laos attracts many wildlife trade tourists simply because wildlife products are more available. Throughout this value chain, people are exposed to blood, viscera, feces, and saliva of rodents, and can be further exposed to these materials during the butchering process, where butchers can accidentally cut themselves with knives, allowing for efficient transmission of viruses from rodents to humans [9]. Considering the significant interactions of wildlife and especially rodents with humans in Laos, we were interested in investigating the presence of CoVs in these animals, which can be primary or intermediate hosts for CoVs with zoonotic potential.

Samples were collected from both live and freshly killed animals, either trapped in or around village homes, or voluntarily provided by local hunters upon their return to the village following hunting forays. With permission of market operators and vendors, samples were also collected from freshly killed animals for sale in markets. Oral and rectal swab specimens were collected in duplicate into individual 1.5 mL screw-top cryotubes containing either 500 μl of Trizol® (Invitrogen), or of Universal Viral Transport Medium (BD) respectively. Samples were then directly placed in liquid nitrogen dry shippers and upon arrival at the laboratory transferred into a −80°C freezer. Staff wore N95 masks, nitrile gloves, dedicated clothing, washable shoes or shoe covers, and protective eyewear during both live and dead animal sampling.

RNA was extracted using the Zymo Direct-zol RNA kit and stored at −80°C until analysis. RNA was converted into cDNA using a Maxima H Minus First Strand cDNA Synthesis Kit (Thermo Scientific) and was stored at −20°C until analysis. Two conventional nested broad range PCR assays, both targeting conserved regions in the RNA-Dependent RNA Polymerase gene (RdRp) were used to test the samples for CoV cDNA. The first PCR amplifies a product of approximately 286nt between the primer binding sites. The first round (CoV-FWD1: CGT TGG IAC WAA YBT VCC WYT ICA RBT RGG and CoV-RVS1: GGT CAT KAT AGC RTC AVM ASW WGC NAC ATG) and second round (CoV-FWD2: GGC WCC WCC HGG NGA RCA ATT and CoV-RVS2: GGW AWC CCC AYT GYT GWA YRT C) primers of this PCR were specifically designed for the detection of a broad range of CoVs [10]. The second PCR was used in two modified versions, one of them specifically targeting a broad range of CoVs in bats the second one broadly targeting CoVs of other hosts. In both cases, the first round of the semi nested PCR utilized the primers CoV-FWD3 (GGT TGG GAY TAY CCH AAR TGT GA) and CoV-RVS3 (CCA TCA TCA SWY RAA TCA TCA TA) for the first round. In the second round either CoV-FWD4/Bat (GAY TAY CCH AAR TGT GAY AGA GC) or CoV-FWD4/Other (GAY TAY CCH AAR TGT GAU MGW GC) were used as forward primers, while the reverse primer was again CoV-RVS3 [11]. Both versions amplify 387nt between the primer binding sites. CoV RNA positive samples were subjected to Cytochrome b PCR to verify the host species. The primers CytB_F (GAG GMC AAA TAT CAT TCT GAG G) and CytB_R (TAG GGC VAG GAC TCC TCC TAG T) were used to amplify a primer-flanked 435nt fragment of the highly conserved mitochondrial gene [12].

PCR products were subjected to gel electrophoresis on a 1.5% agarose gel and products of the expected amplicon sizes were excised. DNA was extracted using the Qiagen QIAquick Gel Extraction Kit and was sent for commercial Sanger sequencing (1st BASE). All results from sequencing were analyzed in the Geneious 7.1 software, primer trimmed, and consensus sequences compared to the GenBank database (BLAST N, NCBI).

All sequences were deposited in GenBank under submission numbers MT083286, MT083287, MT083291-MT083296, MT083363-MT083365 and MT083405.

Maximum likelihood phylogenetic trees were constructed including different genera (Alpha, Beta and Gamma) and species of known CoVs as well as species/sub-species detected in Laos during the PREDICT project. Only a single sequence was included for isolates with nucleotide identities of more than 95%. Multiple sequence alignments were made in Geneious (version 11.1.3, MUSCLE Alignment), and regions supported by less than 50% of the sequences were excluded. Bayesian phylogeny of the polymerase gene fragment was inferred using MrBayes (version 3.2) with the following parameters: Datatype=DNA, Nucmodel=4by4, Nst=1, Coavion=No, # States=4, Rates=Equal, 2 runs, 4 chains of 1,000,000 generations. The sequence of a whale *Gammacoronavirus* served as outgroup to root the trees, and trees were sampled after every 1,000 steps during the process to monitor phylogenetic convergence [13]. The average standard deviation of split frequencies was below 0.0074 for the Watanabe PCR based analysis and below 0.0054 for the Quan PCR based analysis (MrBayes recommended final average <0.01). The first 10% of the trees were discarded and the remaining ones combined using TreeAnnotator (version 2.5.1; http://beast.bio.ed.ac.uk) and displayed with FIGTREE (1.4.4; http://tree.bio.ed.ac.uk/) [14].

During the sampling phases of the project (2010-2013 and 2016-2018) 851 rodents, 124 carnivores, 44 primates, 8 tree shrews and one colugo (*Galeopterus variegatus*) were sampled and tested for the presence of CoV RNA (Figure 1). Rodents belonged to the families Sciuridae (475), Muridae (370), Diatomyidae (5), and Hystricidae (1); carnivores belonged to the families Viverridae (121), Mustelidae (2), and Felidae (1); primates belonged to the families Cercopithecidae (23), Lorisidae (18), and Hylobatidae (2), while one remained unidentified, and all tree shrews were of the Tupaiidae family (8) (Supplement 1).

**Figure 1.**
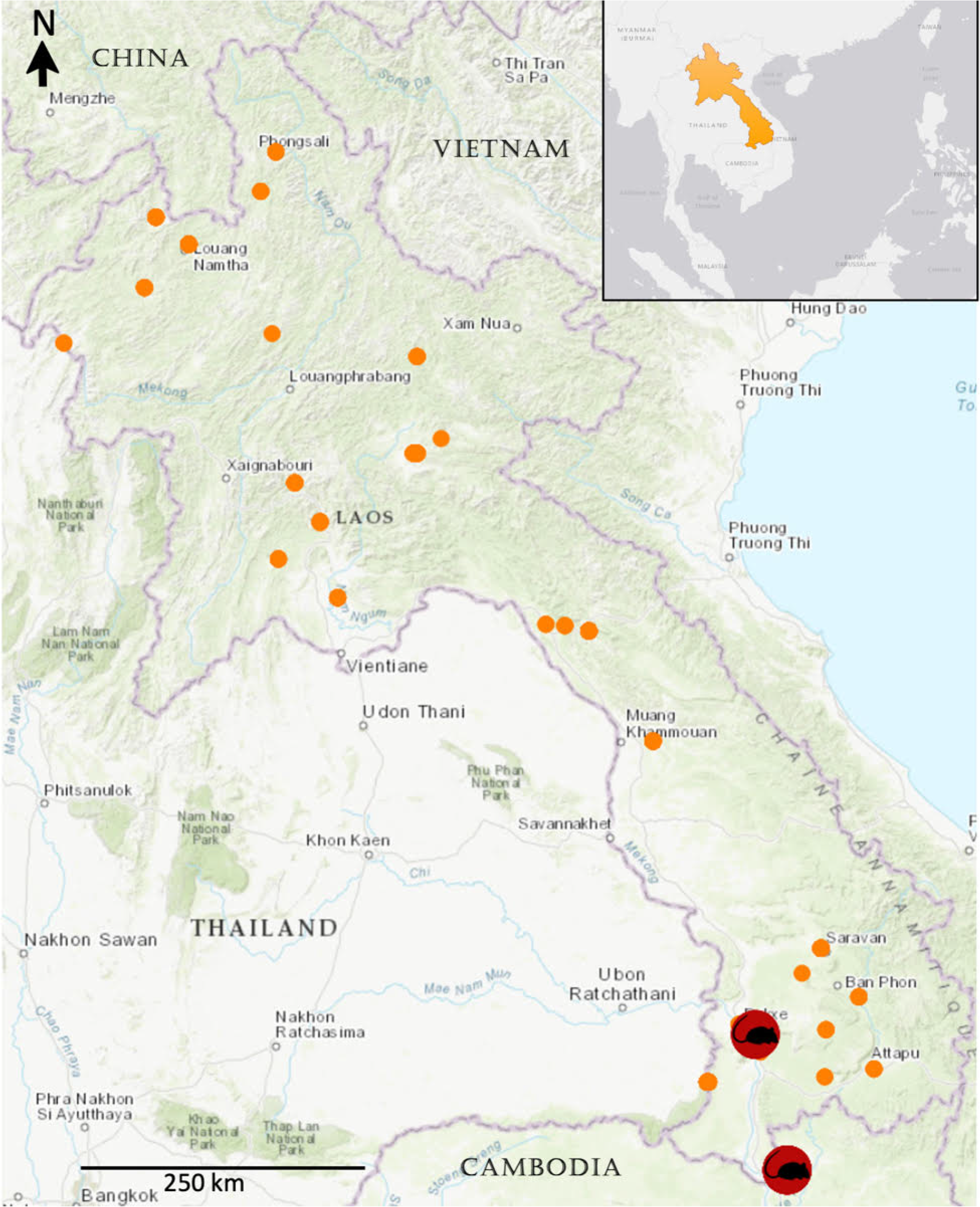
Geographical map indicating all sampling sites within Laos (orange dots), highlighting the locations where coronaviruses in rodents were detected.

CoV RNA was detected in 12 rodents, which corresponds to 1.4% of sampled rodents. All 12 of them were sampled in the south of Laos, and all but one were oral swab samples (Figure 1, Supplement 1). No CoV RNA was detected in any of the carnivores, primates, tree shrews, or the colugo. In all 12 positive animals, it was the Watanabe PCR that amplified CoV nucleic acid. Eleven of the isolates were identical or very similar to each other, and all 12 were similar to CoVs found in different rodents in neighboring China and Vietnam. Nine of the CoV RNA positive animals were caught in the same area within a time window of 5 days (Figure 2, Table 1). Nine of the positive animals, all *Rattus exulans*, were found in and around human dwellings, while the other were three squirrels (one *Dremomys rufigenis*, two *Menetes berdmorei*), sampled at a wet market not far from Pakse, the capital city of Champasak province, where they were being sold for consumption. Eleven of the positive animals were sampled in the dry season (December), and one was sampled in the wet season (June).

**Table 1:** List of CoV RNA positive rodent samples.

| Group | Isolate & Genbank | Host | BLAST N<br>2020-02-25 | Collection date & PCR |
| --- | --- | --- | --- | --- |
| W-Beta-5 | LAAR0140<br>MT083291 | Rodent,<br>Rattus exulans | 97% Coronaviridae sp.<br>isolate 7565L07R6RATCoV<br>(KX092227) | 2016-12-09<br>Watanabe |
| W-Beta-5 | LAAR0156<br>MT083292 | Rodent,<br>Rattus exulans | 97% Coronaviridae sp.<br>isolate 7565L07R6RATCoV<br>(KX092227) | 2016-12-09<br>Watanabe |
| W-Beta-5 | LAAR0174<br>MT083293 | Rodent,<br>Rattus exulans | 97% Coronaviridae sp.<br>isolate 7565L07R6RATCoV<br>(KX092227) | 2016-12-10<br>Watanabe |
| W-Beta-5 | LAAR0198<br>MT083294 | Rodent,<br>Rattus exulans | 97% Coronaviridae sp.<br>isolate 7565L07R6RATCoV<br>(KX092227) | 2016-12-12<br>Watanabe |
| W-Beta-5 | LAAR0202<br>MT083295 | Rodent,<br>Rattus exulans | 97% Coronaviridae sp.<br>isolate 7565L07R6RATCoV<br>(KX092227) | 2016-12-12<br>Watanabe |
| W-Beta-5 | LAAR0206<br>MT083363 | Rodent,<br>Rattus exulans | 97% Coronaviridae sp.<br>isolate 7565L07R6RATCoV<br>(KX092227) | 2016-12-12<br>Watanabe |
| W-Beta-5 | LAAR0208<br>MT083296 | Rodent,<br>Rattus exulans | 97% Coronaviridae sp.<br>isolate 7565L07R6RATCoV<br>(KX092227) | 2016-12-12<br>Watanabe |
| W-Beta-5 | LAAR0209<br>MT083286 | Rodent,<br>Rattus exulans | 97% Coronaviridae sp.<br>isolate 7565L07R6RATCoV<br>(KX092227) | 2016-12-13<br>Watanabe |
| W-Beta-5 | LAAR0210<br>MT083287 | Rodent,<br>Rattus exulans | 97% Coronaviridae sp.<br>isolate 7565L07R6RATCoV<br>(KX092227) | 2016-12-13<br>Watanabe |
| W-Beta-5 | LAAR0235<br>MT083364 | Rodent,<br>Dremomys<br>rufigenis | 97% Coronaviridae sp.<br>isolate 7565L07R6RATCoV<br>(KX092227) | 2016-12-18<br>Watanabe |
| W-Beta-5 | LAAR0239<br>MT083365 | Rodent,<br>Menetes<br>berdmorei | 97% Coronaviridae sp.<br>isolate 7565L07R6RATCoV<br>(KX092227) | 2016-12-19<br>Watanabe |
| W-Beta-6 | LAAR0561<br>MT083405 | Rodent,<br>Menetes<br>berdmorei | 94% Rodent coronavirus<br>isolate RtMc-CoV-2/YN2013<br>(KY370058) | 2018-06-12<br>Watanabe |

**Figure 2.**
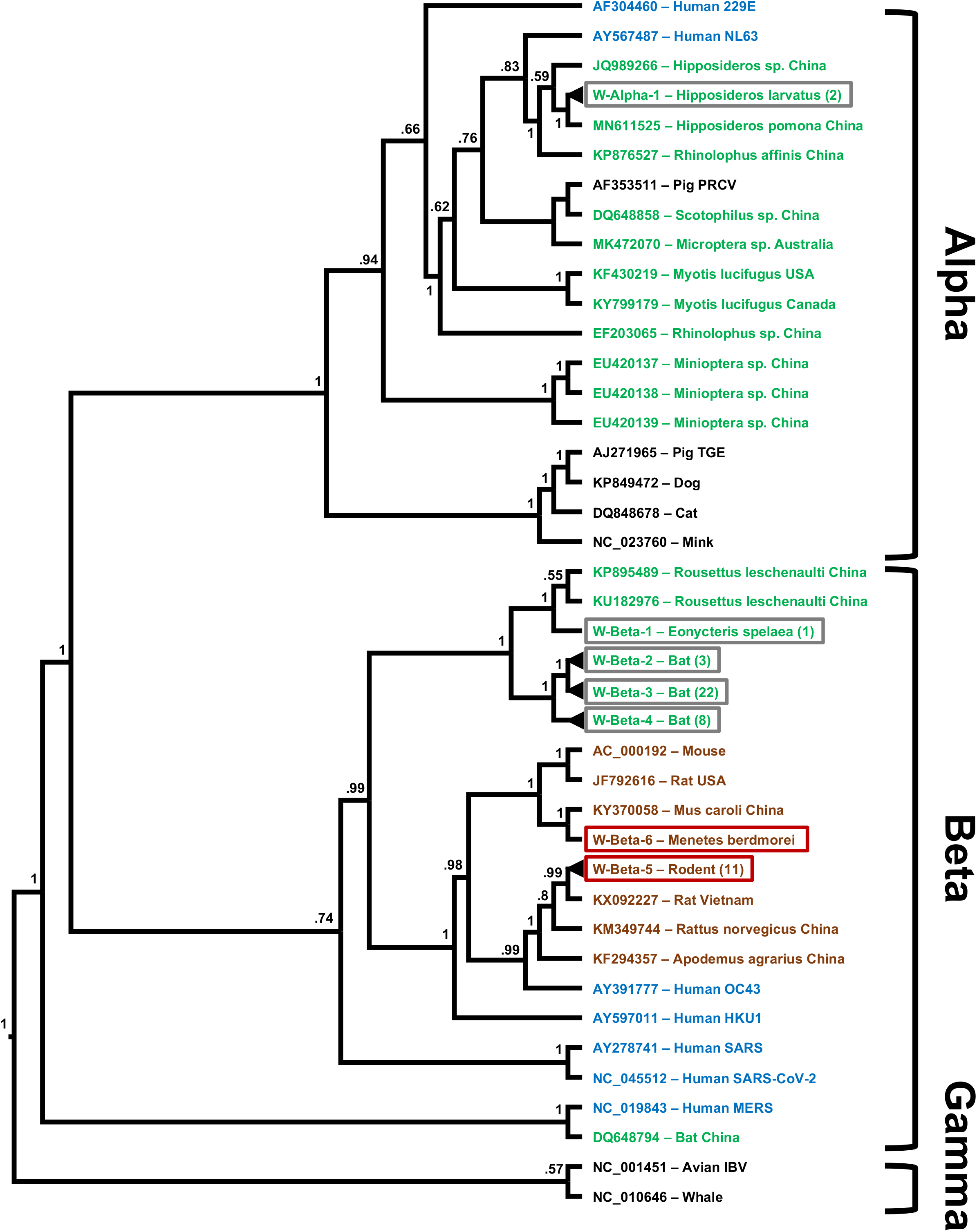
Maximum likelihood phylogenetic tree of coronaviruses presented as a proportional cladogram, based on the RdRp region targeted by the PCR by Watanabe et. al. [11]. The tree includes the sequences detected here (red boxes) and those described previously in Laos (grey boxes) and indicates the number of isolates with less than 5% difference in brackets for isolates. GenBank accession numbers are listed for published sequences from outside of Laos, while sequences obtained during the PREDICT project are identified by cluster names (compare Table 1 and Supplemental 3). Green font indicates coronavirus sequences obtained from bats, brown font indicates rodents, blue humans and black other hosts. The host species and country of sequence origin are indicated for bats and rodents if applicable, no species is indicated for isolates if detected in more than 1 species (compare Table 1 and Supplement 3). Numbers at nodes indicate bootstrap support.

Since there are abundant contact opportunities for wildlife pathogens and humans in Laos, and considering that coronavirus-zoonotic events can involve intermediate hosts, as in the cases of SARS and MERS, we focused our screening on non-bat species potentially capable of playing that role. We found CoV RNA in swab samples from several of the rodents examined in the study. Earlier, we noted a relatively high number of diverse CoVs detected in various species of bats all over Laos (Supplement 2 & 3) [5]. This corresponds to similar findings in other countries with a tropical climate, and a hypothesis has been suggested that bats may be serving as a seeding host for zoonotic CoV infections [15,16]. The 1.4% prevalence of CoV RNA in rodents was much lower than what had been detected in bats in Laos, however such has been observed repeatedly, re-emphasizing the role of bats as a primary CoV source [5,15,16–18]. It is worth noting, that we targeted rodents most likely to be in contact with humans and transmit virus, and did find fewer CoV RNA positive animals than other studies of rodents in the region or elsewhere. A variety of factors may explain the lower incidence of corona virus positive animals in this study including the sample types tested. Studies where CoV RNA was more frequently detected among rodents have used intestine or fecal matter for their studies, while we tested oral and rectal swab samples [18–20]. We employed swab sampling to minimize harm to live animals, and to avoid any damage to rodents possessed by hunters or market vendors during sampling. Organ collection was rarely feasible, even from dead rodents in market settings, since size and mass of the animal are used to determine the selling price. While potentially underestimating the actual CoV circulation, swab sampling has the advantage of being minimally invasive, quick, and applicable to all species.

The 12 rodent CoV sequences we found fall into two clusters, with 11 of them differing by only one nucleotide. Therefore, these 11 likely belong to the same strain that may have been circulating at that time, since they were obtained from rodents in the same southern region of Laos during December 2016 (Table 1). Both CoV strains detected here cluster with other *Betacoronaviruses* previously detected in rodents in the region (Figure 2, Table 1). This suggests that the viruses had a longer evolutionary history within rodent hosts and probably did not derive from a recent cross species transmission event.

None of the CoVs detected in Laos wildlife, neither the ones described earlier in bats nor the ones described here in rodents, have a very close connection to CoVs currently known to cause human disease. The rodent coronaviruses do fall into the same cluster as human coronaviruses OC43 and HKU1 though, which are believed to be derived from a direct or indirect spill-over of rodent viruses to humans [21]. However, we still do not know enough about the molecular mechanisms and drivers of zoonotic events to determine risk or lack of risk with certainty. We conclude that Laos’ wildlife does harbor diverse CoVs, and that a potential for interspecies transmission of viruses and novel diseases exists. Human contact with wildlife like bats and rodents is common throughout the country, with many rural households consuming bushmeat as a main source of protein and utilizing it as a trade commodity, this risk potential is particularly relevant. Therefore, behavioral risk reduction, vigilance, ongoing surveillance and research are important to help mitigate the risks of coronavirus zoonotic disease emergence and transmission in the region, especially in the aftermath of COVID-19.

## Acknowledgements

The authors would like to thank: The government of Laos for the permission to conduct this study; the staff of the National Animal Health Laboratory in Vientiane, Wildlife Conservation Society, and Metabiota who assisted in sample collection and testing, as well as Vibol Hul and Philippe Dussart from the Institut Pasteur du Cambodge; any other involved members of the PREDICT-1 and PREDICT-2 consortium (https://ohi.vetmed.ucdavis.edu/programs-projects/predict-project/authorship). This study was made possible by the generous support of the American people through the United States Agency for International Development (USAID) Emerging Pandemic Threats PREDICT program (cooperative agreement numbers GHN-A-OO-09-00010-00 and AID-OAA-A-14-00102). The contents are the responsibility of the authors and do not necessarily reflect the views of USAID or the United States Government.

## Compliance with Ethical Standards

Animal capture and specimen collection was approved by the Institutional Animal Care and Use Committee (IACUC, UC Davis) and the Ministry of Agriculture and Forestry of Lao PDR. Philippe Buchy is currently an employee of GSK vaccines.

## References

1. Zhu N, Zhang D, Wang W, et al. A Novel Coronavirus from Patients with Pneumonia in China, 2019. N Engl J Med. January 2020:NEJMoa2001017. doi:10.1056/NEJMoa2001017

2. Lu R, Zhao X, Li J, et al. Genomic characterization and epidemiology of 2019 novel coronavirus: implications for virus origins and receptor binding. The Lancet. January 2020:S0140673620302518. doi:10.1016/S0140-6736(20)30251-8

3. Jones KE, Patel NG, Levy MA, Storeygard A, Balk D, Gittleman JL, Daszak P. Global trends in emerging infectious diseases (2008). Nature 451:990–993

4. Hu B, Zeng LP, Yang XL, Ge XY, Zhang W, Li B, Xie JZ, Shen XR, Zhang YZ, Wang N, Luo DS, Zheng XS, Wang MN, Daszak P, Wang LF, Cui J, Shi ZL. Discovery of a rich gene pool of bat SARS-related coronaviruses provides new insights into the origin of SARS coronavirus (2017). PLoS Pathog 13:e1006698

5. Lacroix A, Duong V, Hul V, San S, Davun H, Omaliss K, Chea S, Hassanin A, Theppangna W, Silithammavong S, Khammavong K, Singhalath S, Greatorex Z, Fine AE, Goldstein T, Olson S, Joly DO, Keatts L, Dussart P, Afelt A, Frutos R, Buchy P. Genetic diversity of coronaviruses in bats in Lao PDR and Cambodia (2017). Infect Genet Evol 48:10–18

6. Han BA, Schmidt JP, Bowden SE, Drake JM. Rodent reservoirs of future zoonotic diseases (2015). Proc Natl Acad Sci USA 112, 7039–7044

7. Olival KJ, Hosseini PR, Zambrana-Torrelio C, Ross N, Bogich TL, Daszak P. Host and viral traits predict zoonotic spillover from mammals (2017). Nature 546, 646–650

8. Greatorex ZF, Olson SH, Singhalath S, Silithammavong S, Khammavong K, Fine AE, Weisman W, Douangngeun B, Theppangna W, Keatts L, Gilbert M, Karesh WB, Hansel T, Zimicki S, O’Rourke K, Joly DO, Mazet JA. Wildlife Trade and Human Health in Lao PDR: An Assessment of the Zoonotic Disease Risk in Markets (2016). PLoS One 11:e0150666

9. Pruvot M, Khammavong K, Milavong P, Philavong C, Reinharz D, Mayxay M, Rattanavong S, Horwood P, Dussart P, Douangngeun B, Theppangna W, Fine AE, Olson SH, Robinson M, Newton P. Toward a quantification of risks at the nexus of conservation and health: The case of bushmeat markets in Lao PDR (2019). Sci Total Environ 676:732–745

10. Quan PL, Firth C, Street C, Henriquez JA, Petrosov A, Tashmukhamedova A, Hutchison SK, Egholm M, Osinubi MO, Niezgoda M, Ogunkoya AB, Briese T, Rupprecht CE, Lipkin WI. Identification of a severe acute respiratory syndrome coronavirus-like virus in a leaf-nosed bat in Nigeria (2010). mBio 1 pii: e00208–10

11. Watanabe S, Masangkay JS, Nagata N, Morikawa S, Mizutani T, Fukushi S, Alviola P, Omatsu T, Ueda N, Iha K, Taniguchi S, Fujii H, Tsuda S, Endoh M, Kato K, Tohya Y, Kyuwa S, Yoshikawa Y, Akashi H. Bat coronaviruses and experimental infection of bats, the Philippines (2010). Emerg Infect Dis 2010 16:1217–1223

12. Townzen JS, Brower AV, Judd DD. Identification of mosquito bloodmeals using mitochondrial cytochrome oxidase subunit I and cytochrome b gene sequences (2008). Med. Vet. Entomol. 22:386–93

13. Ronquist F, Teslenko M, van der Mark P, Ayres DL, Darling A, Höhna S, Larget B, Liu L, Suchard MA, Huelsenbeck JP. MRBAYES 3.2: Efficient Bayesian phylogenetic inference and model selection across a large model space (2012). Syst Biol 61:539–542

14. Bouckaert R, Vaughan TG, Barido-Sottani J, Duchêne S, Fourment M, Gavryushkina A, Heled J, Jones G, Kühnert D, De Maio N, Matschiner M, Mendes FK, Müller NF, Ogilvie HA, du Plessis L, Popinga A, Rambaut A, Rasmussen D, Siveroni I, Suchard MA, Wu CH, Xie D, Zhang C, Stadler T, Drummond AJ. BEAST 2.5: An advanced software platform for Bayesian evolutionary analysis (2019). PLoS Comput Biol 15:e1006650

15. Yuen KY, Lau SK, Woo PC. Wild animal surveillance for coronavirus HKU1 and potential variants of other coronaviruses (2012). Hong Kong Med J.18 Suppl 2:25–6

16. Anthony SJ, Johnson CK, Greig DJ, Kramer S, Che X, Wells H, Hicks AL, Joly DO, Wolfe ND, Daszak P, Karesh W, Lipkin WI, Morse SS; PREDICT Consortium, Mazet JAK, Goldstein T. Global patterns in coronavirus diversity (2017). Virus Evol 3:vex012

17. Wang W, Lin XD, Guo WP, Zhou RH, Wang MR, Wang CQ, Ge S, Mei SH, Li MH, Shi M, Holmes EC, Zhang YZ. Discovery, diversity and evolution of novel coronaviruses sampled from rodents in China (2015). Virology 474:19–27

18. Berto A, Anh PH, Carrique-Mas JJ, Simmonds P, Van Cuong N, Tue NT, Van Dung N, Woolhouse ME, Smith I, Marsh GA, Bryant JE, Thwaites GE, Baker S, Rabaa MA; VIZIONS consortium. Detection of potentially novel paramyxovirus and coronavirus viral RNA in bats and rats in the Mekong Delta region of southern Viet Nam. Zoonoses Public Health 65:30–42

19. Ge XY, Yang WH, Zhou JH, Li B, Zhang W, Shi ZL, Zhang YZ. Detection of alpha- and betacoronaviruses in rodents from Yunnan, China (2017). Virol J 14:98

20. Monchatre-Leroy E, Boué F, Boucher JM, Renault C, Moutou F, Ar Gouilh M, Umhang G. Identification of Alpha and Beta Coronavirus in Wildlife Species in France: Bats, Rodents, Rabbits, and Hedgehogs (2017). Viruses 9. pii: E364

21. Cui J, Li F, Shi ZL, Origin and evolution of pathogenic coronaviruses (2019). Nat Rev Microbiol 17:181–192

