## Supplement 2 for "Coronavirus surveillance of wildlife in the Lao People’s Democratic Republic detects viral RNA in rodents"

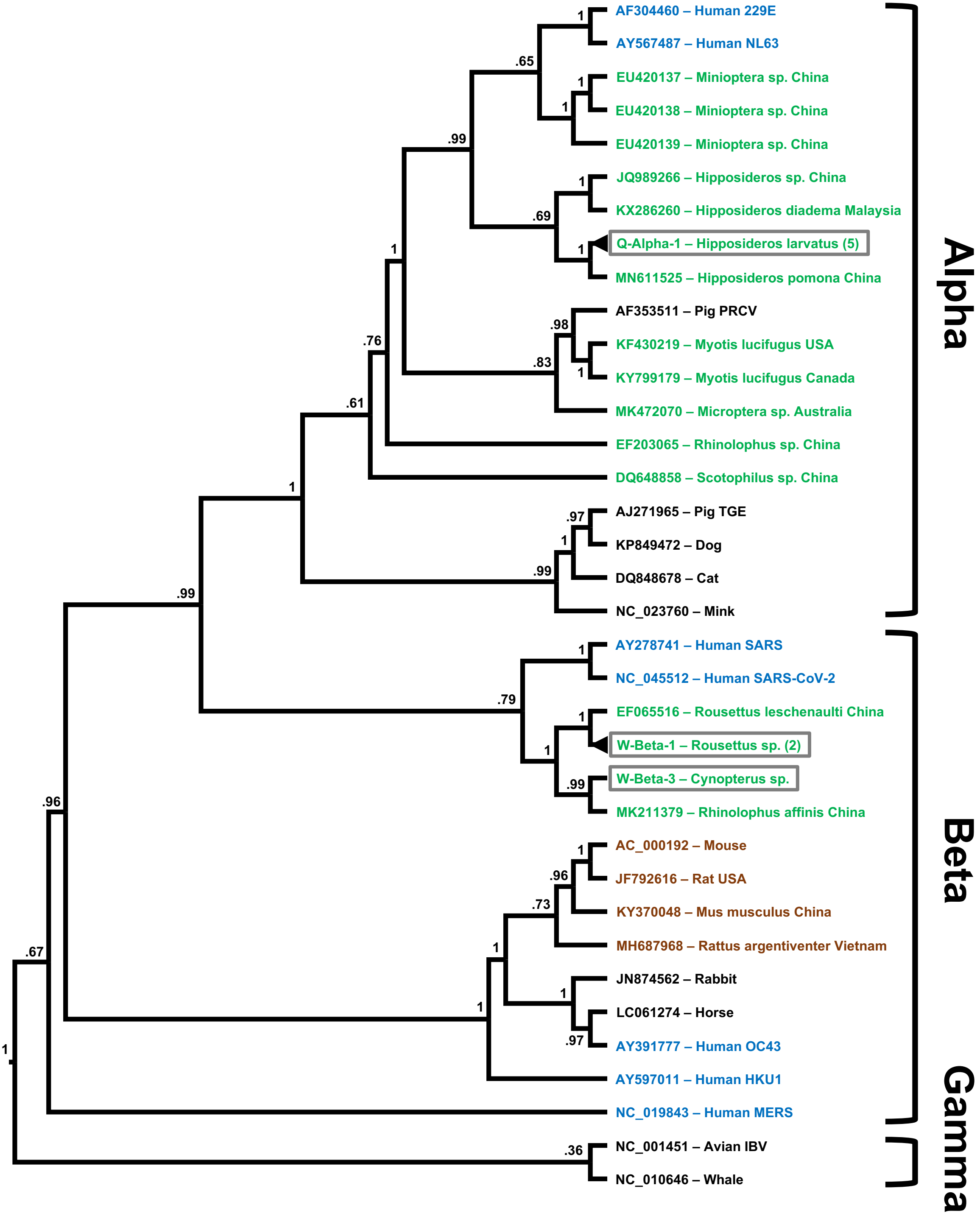

AF304460 – Human 229E
AY567487 – Human NL63
EU420137 – Miniopterа sp. China
EU420138 – Miniopterа sp. China
EU420139 – Miniopterа sp. China
JQ989266 – Hipposideros sp. China
KX286260 – Hipposideros diadema Malaysia
Q-Alpha-1 – Hipposideros larvatus (5)
MN611525 – Hipposideros pomona China
AF353511 – Pig PRCV
KF430219 – Myotis lucifugus USA
KY799179 – Myotis lucifugus Canada
MK472070 – Microptera sp. Australia
EF203065 – Rhinolophus sp. China
DQ648858 – Scotophilus sp. China
AJ271965 – Pig TGE
KP849472 – Dog
DQ848678 – Cat
NC_023760 – Mink
AY278741 – Human SARS
NC_045512 – Human SARS-CoV-2
EF065516 – Rousettus leschenaulti China
W-Beta-1 – Rousettus sp. (2)
W-Beta-3 – Cynopterus sp.
MK211379 – Rhinolophus affinis China
AC_000192 – Mouse
JF792616 – Rat USA
KY370048 – Mus musculus China
MH687968 – Rattus argentiventer Vietnam
JN874562 – Rabbit
LC061274 – Horse
AY391777 – Human OC43
AY597011 – Human HKU1
NC_019843 – Human MERS
NC_001451 – Avian IBV
NC_010646 – Whale
Alpha
Beta
Gamma
1
.65
1
1
.99
1
.69
1
1
.98
.83
1
.76
.61
1
.97
1
.99
.99
1
.79
1
1
.99
.96
1
.96
.73
1
1
1
.97
.67
1
.36

**Legend: Supplement 2**

Maximum likelihood phylogenetic tree of coronaviruses presented as a proportional cladogram, based on the RdRp region targeted by the PCR by Quan et. al. [10]. The tree includes sequences of bat coronaviruses detected in Laos (grey boxes) and indicates the number of isolates with less than 5% difference in brackets [5]. GenBank accession numbers are listed for previously published sequences, while sequences obtained during the project are identified by cluster names (Supplemental 3). Green font indicates coronavirus sequences obtained from bats, brown font indicates rodents, blue humans and black other hosts. The host species and country of sequence origin are indicated for bats and rodents if applicable, no species is indicated for novel isolates if detected in more than 1 species (Supplemental 3). Numbers at nodes indicate bootstrap support.
