## Supplement 3 for "Coronavirus surveillance of wildlife in the Lao People’s Democratic Republic detects viral RNA in rodents"

**Supplement 3: Table of CoV RNA positive bat samples [5]**

| Group | Isolate | Host | BLAST N 2020-02-25 | Year & PCR |
|---|---|---|---|---|
| W-Beta-3 | LAP11-A0058 KX284902 | Bat, Rousettus sp. | 99% Rousettus bat coronavirus HKU9 clone 2166 (MG762620) | 2011 Watanabe |
| W-Beta-3 | LAP11-A0071 KX284903 | Bat, Rousettus sp. | 99% Rousettus bat coronavirus HKU9 PREDICT-KHP13-BN1-0021 (KX285761) | 2011 Watanabe |
| W-Beta-3 | LAP11-A0077 KX284904 | Bat, Rousettus sp. | 99% Rousettus bat coronavirus HKU9 clone 9466 (MG762663) | 2011 Watanabe |
| W-Beta-3 | LAP11-A0094 KX284905 | Bat, Rousettus sp. | 99% Rousettus bat coronavirus HKU9 clone 162387 (MG762664) | 2011 Watanabe |
| W-Beta-1 | LAP11-D0063 KX284906 | Bat, Eonycteris spelaea | 99% Bat coronavirus isolate ML42C (KU182976) | 2011 Watanabe |
| W-Beta-3 | LAP11-D0064 KX284907 | Bat, Rousettus sp. | 100% Rousettus bat coronavirus HKU9 clone 9466 (MG762663) | 2011 Watanabe |
| W-Beta-3 | LAP11-J0087 KX284908 | Bat, Eonycteris spelaea | 100% Rousettus bat coronavirus HKU9 clone 9466 (MG762663) | 2011 Watanabe |
| W-Beta-3 | LAP11-J0091 KX284909 | Bat, Rousettus leschenaultii | 99% Rousettus bat coronavirus HKU9 clone 9466 (MG762663) | 2011 Watanabe |
| W-Beta-3 | LAP11-J0095 KX284910 | Bat, Rousettus amplexicaudatus | 100% Rousettus bat coronavirus HKU9 clone 9466 (MG762663) | 2011 Watanabe |
| W-Beta-2 | LAP11-K0006 KX284911 | Bat, Eonycteris spelaea | 99% Rousettus bat coronavirus HKU9 clone 2171 (MG762621) | 2011 Watanabe |
| W-Beta-2 | LAP11-K0012 KX284912 | Bat, Eonycteris spelaea | 99% Rousettus bat coronavirus HKU9 clone 2171 (MG762621) | 2011 Watanabe |
| W-Beta-2 | LAP11-K0038 KX284913 | Bat, Rousettus amplexicaudatus | 99% Rousettus bat coronavirus HKU9 clone 2171 (MG762621) | 2011 Watanabe |
| W-Beta-3 | LAP11-K0040 KX284914 | Bat, Rousettus amplexicaudatus | 99% Rousettus bat coronavirus HKU9 PREDICT-KHP13-BN1-0021 (KX285761) | 2011 Watanabe |
| W-Beta-3 | LAP11-M0052 | Bat, Rousettus sp. | 99% Rousettus bat coronavirus HKU9 clone | 2011 Watanabe |

| | | | | |
|---|---|---|---|---|
| | KX284915 | | 162387 (MG762664) | |
| W-Beta-4 | LAP11-M0053 KX284916 | Bat, Rousettus sp. | 100% Rousettus bat coronavirus HKU9 clone 9433 (MG762660) | 2011 Watanabe |
| W-Beta-3 | LAP11-M0054 KX284917 | Bat, Rousettus sp. | 99% Rousettus bat coronavirus HKU9 clone 2166 (MG762620) | 2011 Watanabe |
| W-Beta-4 | LAP12-A0005 KX284918 | Bat, Rousettus sp. | 100% Rousettus bat coronavirus HKU9 isolate Rousettus spp / Jinghong / 2009 (MG762674) | 2012 Watanabe |
| W-Beta-3 | LAP12-A0008 KX284919 | Bat, Eonycteris spelaea | 99% Rousettus bat coronavirus HKU9 clone 9466 (MG762663) | 2012 Watanabe |
| W-Beta-3 | LAP12-A0010 KX284920 | Bat, Rousettus sp. | 99% Rousettus bat coronavirus HKU9 clone 9466 (MG762663) | 2012 Watanabe |
| W-Beta-4 | LAP12-A0011 KX284921 | Bat, Rousettus sp. | 99% Rousettus bat coronavirus HKU9 clone 9433 (MG762660) | 2012 Watanabe |
| W-Beta-4 | LAP12-A0012 KX284922 | Bat, Rousettus sp. | 100% Rousettus bat coronavirus HKU9 isolate Rousettus spp / Jinghong / 2009 (MG762674) | 2012 Watanabe |
| W-Beta-4 | LAP12-A0019 KX284923 | Bat, Rousettus sp. | 99% Rousettus bat coronavirus HKU9 isolate Rousettus spp / Jinghong / 2009 (MG762674) | 2012 Watanabe |
| W-Beta-3 | LAP12-A0021 KX284924 | Bat, Rousettus sp. | 99% Rousettus bat coronavirus HKU9 clone 9466 (MG762663) | 2012 Watanabe |
| W-Beta-3 | LAP12-A0022 KX284925 | Bat, Rousettus sp. | 99% Rousettus bat coronavirus HKU9 clone 9466 (MG762663) | 2012 Watanabe |
| W-Beta-4 | LAP12-A0024 KX284926 | Bat, Rousettus sp. | 99% Rousettus bat coronavirus HKU9 clone 9433 (MG762660) | 2012 Watanabe |
| W-Beta-3 | LAP12-A1-0007 KX285065 | Bat, Rousettus sp. | 99% Rousettus bat coronavirus HKU9 clone 9466 (MG762663) | 2012 Watanabe |
| Q-Beta-1 | LAP12-A1-0007 KX286293 | Bat, Rousettus sp. | 92% Bat coronavirus HKU9-4 (EF065516) | 2012 Quan |
| W-Beta-3 | LAP12-A1-0014 KX285066 | Bat, Rousettus sp. | 100% Rousettus bat coronavirus HKU9 clone 9466 (MG762663) | 2012 Watanabe |
| Q-Beta- | LAP12- | Bat, | 91% Bat coronavirus HKU9- | 2012 |

| | | | | |
|---|---|---|---|---|
| 1 | A3-0001 KX286294 | Rousettus sp. | 4 (EF065516) | Quan |
| W-Beta-3 | LAP12-D0077 KX284931 | Bat, Rousettus sp. | 99% Rousettus bat coronavirus HKU9 clone 162387 (MG762664) | 2012 Watanabe |
| W-Beta-3 | LAP12-E0004 KX284932 | Bat, Eonycteris spelaea | 99% Rousettus bat coronavirus HKU9 clone 9466 (MG762663) | 2012 Watanabe |
| W-Beta-3 | LAP12-E0016 KX284933 | Bat, Rousettus sp. | 100% Rousettus bat coronavirus HKU9 clone 9466 (MG762663) | 2012 Watanabe |
| W-Beta-4 | LAP12-E0019 KX284934 | Bat, Rousettus sp. | 99% Rousettus bat coronavirus HKU9 isolate Rousettus spp / Jinghong / 2009 (MG762674) | 2012 Watanabe |
| W-Beta-3 | LAP12-E0072 KX284935 | Bat, Rousettus sp. | 99% Rousettus bat coronavirus HKU9 clone 162387 (MG762664) | 2012 Watanabe |
| W-Alpha-1 | LAP12-E1-0041 KX284936 | Bat, Hipposideros larvatus | 89% Hipposideros pomona bat coronavirus CHB25 isolate CHB0025 (MN611525) | 2012 Watanabe |
| Q-Alpha-1 | LAP12-E1-0041 KX284291 | Bat, Hipposideros larvatus | 91% Hipposideros pomona bat coronavirus CHB25 isolate CHB0025 (MN611525) | 2012 Quan |
| Q-Alpha-1 | LAP12-E1-0047 KX284292 | Bat, Hipposideros larvatus | 91% Hipposideros pomona bat coronavirus CHB25 isolate CHB0025 (MN611525) | 2012 Quan |
| W-Alpha-1 | LAP12-E1-0052 KX284937 | Bat, Hipposideros larvatus | 89% Hipposideros pomona bat coronavirus CHB25 isolate CHB0025 (MN611525) | 2012 Watanabe |
| Q-Alpha-1 | LAP12-E1-0052 KX286287 | Bat, Hipposideros larvatus | 91% Hipposideros pomona bat coronavirus CHB25 isolate CHB0025 (MN611525) | 2012 Quan |
| Q-Alpha-1 | LAP12-E1-0058 KX286288 | Bat, Hipposideros larvatus | 91% Hipposideros pomona bat coronavirus CHB25 isolate CHB0025 (MN611525) | 2012 Quan |
| Q-Alpha-1 | LAP12-E1-0063 KX286289 | Bat, Hipposideros larvatus | 91% Hipposideros pomona bat coronavirus CHB25 isolate CHB0025 (MN611525) | 2012 Quan |
| Q-Beta- | LAP12- | Bat, | 99% Coronavirus BtRt- | 2012 |

| | | | | |
|---|---|---|---|---|
| 2 | G1-0029 KX286290 | Cynopterus sp. | BetaCoV/GX2018 (MK211379) | Quan |
| W-Beta-3 | LAP12-I0007 KX284938 | Bat, Rousettus sp. | 99% Rousettus bat coronavirus HKU9 clone 9466 (MG762663) | 2012 Watanabe |
| W-Beta-4 | LAP13-D0039 KX285067 | Bat, Rousettus sp. | 99% Rousettus bat coronavirus HKU9 clone 9433 (MG762660) | 2013 Watanabe |
